# Syntaxin-1 is necessary for UNC5/Netrin-1-dependent macropinocytosis and chemorepulsion

**DOI:** 10.1101/2021.05.20.444954

**Authors:** Ramón Martínez-Mármol, Ashraf Muhaisen, Tiziana Cotrufo, Cristina Roselló-Busquets, Marc Hernaiz-Llorens, Francesc Pérez-Branguli, Rosa Maria Andrés, Oriol Ros, Marta Pascual, Fausto Ulloa, Eduardo Soriano

## Abstract

Brain connectivity requires correct axonal guidance to drive axons to their appropriate targets. This process is orchestrated by guidance cues that exert attraction or repulsion to developing axons. However, the intricacies of the cellular machinery responsible for the correct response of growth cones are just being unveiled. Netrin-1 is a bifunctional molecule involved in axon pathfinding and cell migration that induces repulsion during postnatal cerebellar development. This process is mediated by Uncoordinated locomotion 5 (UNC5) receptors located on external granule layer (EGL) tracts. Here, we demonstrate that this response is characterized by enhanced membrane internalization through macropinocytosis, but not clathrin-mediated endocytosis. We show that UNC5 receptors form a protein complex with the t-SNARE syntaxin-1 (Stx1). By combining botulinum neurotoxins, a shRNA knock-down strategy and Stx1 knock-out mice, we demonstrate that this SNARE protein is required for Netrin-1-induced macropinocytosis and chemorepulsion, suggesting that Stx1 is crucial in regulating Netrin-1-mediated axonal guidance.

## Introduction

During the development of the nervous system, migrating cells and axonal growth cones respond to attractive and repellent guidance cues. Netrin-1 belongs to a family of laminin-related secreted proteins that act as bifunctional guidance factors, generating either chemoattractive or chemorepulsive responses. The response to Netrin-1 is determined by the composition of receptors on the surface of the growth cones. Netrin-1-induced axon attraction is mediated by Deleted in Colorectal Cancer (DCC) receptors (Keino-Masu *et al*, 1996), Neogenin/DCC like molecule (De Vries & Cooper, 2008) and Down Syndrome Cell Adhesion Molecule (DSCAM) receptors (Ly *et al*, 2008). The Uncordinated locomotion 5 family of receptors (UNC5A, B, C and D) mediates repulsion (Keleman & Dickson, 2001). The repulsive action of Netrin-1 requires axonal expression of either UNC5 alone or a complex of UNC5 and either DCC or DSCAM receptors (Hong *et al*, 1999; Purohit *et al*, 2012). Earlier reports described how the expression of a combination of both UNC5 and DCC receptors mediated long-range repulsion in the presence of low concentrations of Netrin-1, whereas UNC5 alone was sufficient to mediate short-range repulsion in the presence of high concentrations of Netrin-1 (Kennedy, 2000). However, more recent studies have pointed to a combinatorial and incremental relationship between both receptors in controlling a particular axon response to the cue (Boyer & Gupton, 2018; Muramatsu *et al*, 2010).

Once axons reach their final destination, neuronal communication starts by releasing neurotransmitters contained in synaptic vesicles. Synaptic vesicle exocytosis occurs through the assembly of the SNARE (Soluble NSF-Attachment protein Receptors) protein complex and requires association of the plasma membrane t-SNAREs syntaxin-1 (Stx1) and SNAP25 with the vesicle v-SNARE VAMP2/Synaptobrevin2 (Jahn & Scheller, 2006). In addition, other synaptic proteins, including Synaptotagmins, Munc-18 and Complexin regulate the exocytotic cycle of synaptic vesicles (Brunger *et al*, 2019). Interestingly, many proteins involved in synaptic exocytosis and endocytosis are highly expressed during development and are enriched in growth cones (Urbina & Gupton, 2020). In fact, a number of studies have supported the notion that axonal guidance mediated by extracellular cues may rely on the control of exocytotic and endocytic events at selected regions of the growth cone (Cotrufo *et al*, 2011; Hines *et al*, 2010; Tojima *et al*, 2007; Tojima *et al*, 2010; Zylbersztejn *et al*, 2012).

During chemoattraction, Netrin-1 triggers the recruitment of its receptor DCC to the plasma membrane and increases growth cone exocytosis (Bouchard *et al*, 2008; Bouchard *et al*, 2004; Matsumoto & Nagashima, 2010). Moreover, Stx1 directly interacts with DCC and is required for Netrin-1-dependent chemoattraction of axons and migrating neurons, both *in vitro* and *in vivo* (Barrecheguren *et al*, 2017; Cotrufo *et al*, 2012; Cotrufo *et al*., 2011). Together these data indicate a tight cross-talk between chemoattractive guidance cue signalling pathways and proteins that regulate exocytosis within growth cones.

In contrast, the relationship between chemorepellent cues and proteins that regulate membrane turnover is less well understood. It has been shown that clathrin-dependent endocytosis drives repulsive growth cone responses to Semaphorin 3A (Tojima *et al*., 2010). It is also known that Ephrin-A2, Sonic hedgehog (Shh) and Slit2 stimulate in dorsal root ganglion, retinal ganglion cell and commissural growth cones, a specific type of clathrin-independent endocytosis of large structures known as macropynocitosis (Guo *et al*, 2012; Jurney *et al*, 2002; Kabayama *et al*, 2009; Ros *et al*, 2018). Interestingly, the vesicle SNARE VAMP-2 mediates the sorting of Neuropilin-1/Plexin-A1 receptors, which is required for repulsion by Semaphorin 3A (Zylbersztejn *et al*., 2012). These studies have led to the assumption that membrane remodelling, acting in coordination with F-actin cytoskeletal reorganization, may contribute to growth cone steering by chemorepellent guidance cues (Gallo & Letourneau, 2004; Piper *et al*, 2006). Nevertheless, the mechanistic role of SNARE proteins in these processes remains unclear.

Here we investigate whether Netrin-1-dependent growth cone collapse and chemorepulsion are associated with enhanced membrane internalization. We show that both Netrin-1-induced collapse and axon chemorepulsion are associated with macropynocytosis. Furthermore, we demonstrate for the first time that the SNARE protein Stx1 co-associates with the Netrin-1 receptors UNC5, and that Stx1 function is required for Netrin-1-induced membrane macropinocytosis and repulsion. Overall, our results underscore a novel Stx1-dependent signalling pathway necessary for membrane internalization in Netrin-1-mediated chemorepulsion.

## Results

### UNC5 receptors co-associate with Stx1 both *in vitro and in vivo*

We have previously demonstrated that the t-SNARE Stx1 interacts with the guidance receptor DCC (Cotrufo *et al*., 2012; Cotrufo *et al*., 2011) and the tropomyosin-related kinase (TrkB) receptor (Fuschini *et al*, 2018), with these associations being necessary for the Netrin-1-dependent chemoattraction and neurotrophin-dependent outgrowth of axons, respectively. These results indicate that Stx1 regulates exocytosis and membrane retrieval in growth cones, and is required for Netrin-1-induced chemotropic guidance *in vitro* and *in vivo* (Ros *et al*, 2015). On the other hand, it has also been suggested that SNARE proteins can be involved in certain types of endocytosis within synapses (Xu *et al*, 2013; Zhang *et al*, 2013), and that some SNAREs interact with proteins involved in endocytosis (Galas *et al*, 2000; Koo *et al*, 2011; Miller *et al*, 2011). In addition, we have found that Stx1 is required for the repulsion of commissural axons (Ros *et al*., 2018). Here, we first investigated whether Stx1 interacts with the chemorepulsive Netrin-1 receptors UNC5B and UNC5C, which are important in the control of the repulsive response of axonal tracts during cerebellar development (Alcantara *et al*, 2000). First, we tested this interaction using an *in vitro* system in which HEK-293T cells were co-transfected with Stx1A or Stx1A-EGFP, together with UNC5B-myc, and tested for co-immunoprecipitation (**Figure 1**). Our results revealed that immunoprecipitation with anti-myc antibodies led to Stx1 association (**Figure 1A**). The reverse immunoprecipitation with anti-Stx1 antibodies also yielded myc-tagged UNC5B (**Figure 1A**). This interaction was maintained using anti-GFP antibodies to pull down Stx1A-EGFP (**Figure 1B**). In similar co-immunoprecipitation experiments, cotransfecting UNC5A-myc or UNC5C-myc with Stx1A-EGFP revealed that Stx1 also co-associates with UNC5A and UNC5C receptors (**Figure 1C**). Next, we asked whether Stx1 ia associated with UNC5 receptors *in vivo*. Using cultured postnatal cerebellar EGL neurons, we found that both proteins partially colocalize in growth cones (**Figure 1D**). Moreover, immunoprecipitation of postnatal day 4 (P4) cerebellar tissue with Stx1 revealed interaction with UNC5C (**Figure 1E**). Finally, to investigate whether Stx1 interacts with other receptors that mediate chemorepulsion, we cotransfected HEK-293T cells with Stx1A-EGFP and Neuropilin-1-HA or Plexin-A1-VSV, the receptors for class III Semaphorins (Gil & Del Rio, 2019). Our results showed that Stx1 does not interact with these Semaphorin receptors (**Figure EV1**).

**Figure 1.**
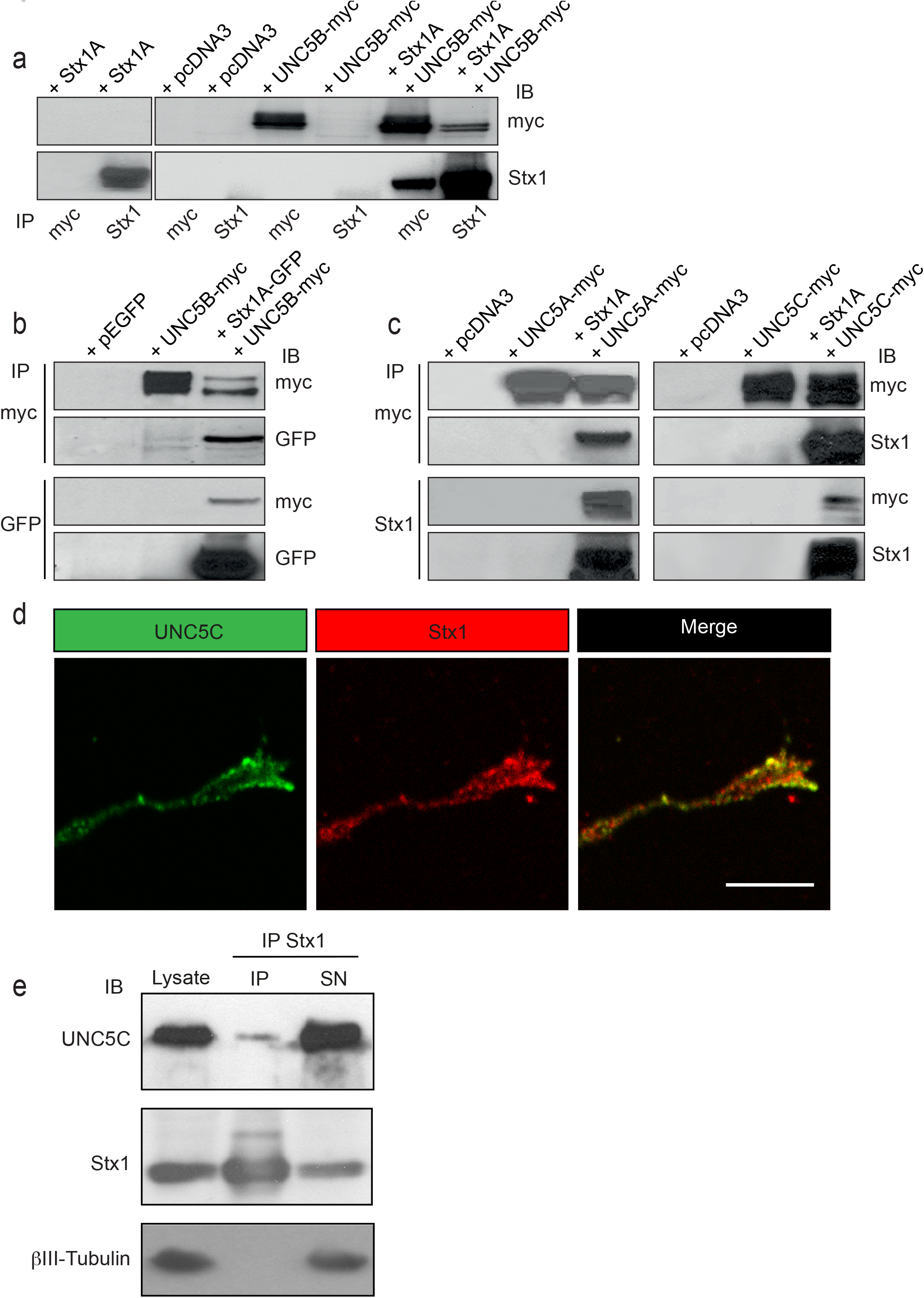
Stx1 interacts with UNC5 receptors. **(A)** HEK-293T cells were transfected with the indicated combination of plasmids (empty vector pcDNA3, Stx1A, UNC5B-myc, or both Stx1A and UNC5B-myc). Protein lysates were immunoprecipitated (IP) with anti-Stx1 or anti-myc antibodies. Co-immunoprecipitation of Stx1A and UNC5B was detected by immunoblotting (IB) using anti-Stx1 or anti-myc antibodies. (**B)** HEK-293T cells were transfected with the indicated combination of plasmids (empty vector pEGFP, Stx1A-GFP, UNC5B-myc, or both Stx1A-GFP and UNC5B-myc). Protein lysates were immunoprecipitated with anti-GFP or anti-myc antibodies. Co-immunoprecipitation of Stx1A and UNC5B was detected by immunoblotting using anti-GFP or anti-myc antibodies. (**C)** HEK-293T cells were transfected with the indicated combination of plasmids (empty vector pcDNA3, Stx1A, UNC5A-myc, UNC5C-myc, Stx1A and UNC5A-myc, or Stx1A and UNC5C-myc). Protein lysates were immunoprecipitated with anti-Stx1 or anti-myc antibodies. Co-immunoprecipitation of Stx1A and UNC5A or UNC5C was detected by immunoblotting using anti-Stx1 or anti-myc antibodies. (**D)** Representative confocal image of an EGL growth cone immunostained against UNC5C and Stx1. Scale bar represents 10 µm. (**E)** UNC5C immunoprecipitated with Stx1 from P4 cerebellar homogenates. Immunoblotting using anti-Stx1 antibody was used as a positive control, and anti-βIII-tubulin antibody was used as a loading control and as a negative immunoprecipitation control.

### Botulinum neurotoxin C1 decreases Netrin-1-dependent chemorepulsion of EGL axons

Netrin-1 exerts long-range chemorepulsion that is important for axonal guidance (Round & Stein, 2007), and plays a major role in the organization of EGL neurons during early postnatal development of the cerebellum, controlling the growth of parallel fibres (Alcantara *et al*., 2000; Hernaiz-Llorens *et al*, 2020; Przyborski *et al*, 1998). EGL neurons have been shown to co-express DCC, neogenin, DSCAM, UNC5B and UNC5C that sense Netrin-1 expressed in the EGL and in interneurons of the molecular layer of the cerebellum (Alcantara *et al*., 2000; Hernaiz-Llorens *et al*., 2020). To investigate whether Stx1 is necessary for Netrin-1-dependent repulsion, we co-cultured postnatal EGL explants in type I collagen 3D hydrogels, and exposed them to aggregates of either control HEK-293T cells or Netrin-1-expressing HEK-293T cells for 2 days (Alcantara *et al*., 2000; Hernaiz-Llorens *et al*., 2020). EGL explants cultured with control HEK-293T cells exhibited a radial pattern of axonal growth (**Figure 2A**), whereas those confronted with Netrin-1-expressing cells exhibited strong chemorepulsion, with most axons growing in the distal quadrant (**Figure 2A**). Botulinum neurotoxins (BoNTs) are metalloproteases that reduce synaptic vesicle exocytosis and neurotransmitter release by cleaving specific SNARE proteins (Schiavo *et al*, 2000). Botulinum neurotoxin A (BoNT/A) exclusively cleaves SNAP25, whereas botulinum neurotoxin C1 (BoNT/C1) cleaves both Stx1 and SNAP25 (Blasi *et al*, 1993; Schiavo *et al*., 2000). Cleavage of Stx1 by BoNT/C1 in EGL cultures was confirmed by western blot (**Figure EV2A**). It has been shown that BoNTs cleaves SNARE proteins in neuronal explants without affecting the secretion of Netrin-1 from stably transfected HEK-293T cells (Cotrufo *et al*., 2011). EGL-derived explants co-cultured with Netrin-1 in the presence of BoNT/C1 or BoNT/A displayed different phenotypes. While Netrin-1 elicited chemorepulsion in EGL explants incubated with BoNT/A, treatment with BoNT/C1 resulted in a radial axonal sprouting (**Figure 2A, B**). As Stx1 interacts with UNC5 receptors, but not with Neuropilin or Plexin receptors, we next investigated whether BoNTs altered Semaphorin 3A/3F-induced chemorepulsion (**Figure EV2B**). Hippocampal explants exhibited strong repulsion when confronted with Semaphorin 3A- or Semaphorin 3F-expressing cells. This repulsion was not altered when the explants were cultured in the presence of BoNT/A or BoNT/C1 (**Figure EV2B**). Taken together, these data indicate that the cleavage of Stx1 specifically abolishes Netrin-1-induced repulsion, but not class III Semaphorin-induced chemorepulsion.

**Figure 2.**
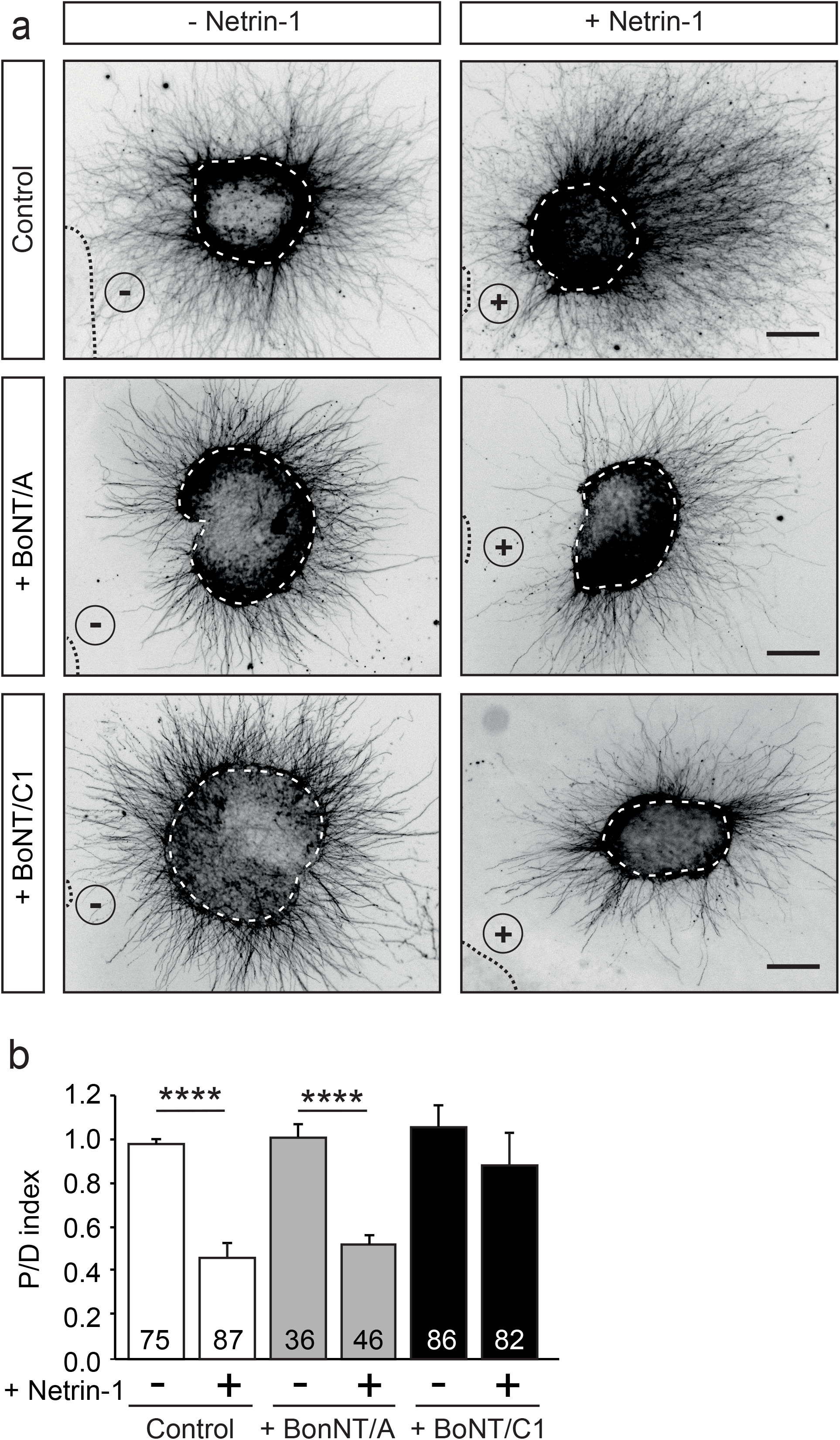
Cleavage of Stx1 by BoNT/C1 blocks Netrin-1-induced repulsion in EGL explants. **(A)** Representative images of EGL explants from P4 mice, immunodetected with anti-βIII-tubulin. Explants were confronted with control cell aggregates (-Netrin-1) or Netrin-1-secreting aggregates (+ Netrin-1). Explants were cultured in the absence of BoNTs (control) or in medium supplemented with 25 nM BoNT/A (+ BoNT/A) or 15 nM BoNT/C1 (BoNT/C1). HEK-293T aggregates are outlined with a dashed line. BoNT/C1, but not BoNT/A, cleaves Stx1 thereby abolishing the netrin-induced repulsion. Scale bar represents 100 µm. (**B)** Graph showing the calculated P/D ratio. Data represent mean ± SEM. One-way ANOVA followed by Dunn’s multiple comparison post hoc test of selected pairs was used. ****p ≤ 0.0001. The number of explants analysed for each condition is indicated within each bar. The results were generated from at least three independent experiments.

### Postnatal Stx1B knock-out mice display decreased Netrin-1 chemorepulsion

Stx1 consists of two similar paralogs in mammals, Stx1A and Stx1B (Gerber *et al*, 2008; Ros *et al*., 2018). We investigated the expression of Stx1A and Stx1B in P3-5 postnatal cerebellum by western blot and found that, although both paralogs are expressed, the postnatal cerebellum shows a preponderant expression of Stx1B (**Figure 3A, B**). It has previously been shown that mice deficient for Stx1A do not display anatomical abnormalities and only exhibit minor physiological neurotransmitter release deficits, most likely due to functional redundancy with Stx1B (Fujiwara *et al*, 2006). To evaluate the importance of Stx1B in Netrin-1-dependent chemorepulsion, we used Stx1B knock-out mice (Ros *et al*., 2018) . The level of Stx1B paralog was specifically depleted in the knock-out mice (-/-), compared with the wild-type (+/+) or the heterozygous (+/-) animals **(Figure EV3A**). Newborn homozygous Stx1B knock-out mice were slightly smaller than their control wild-type littermates (**Figure EV3B**) and exhibited obvious motor deficits which became more apparent with increasing age. Whereas homozygous Stx1B knock-out mice usually survived until P7-P15, heterozygous Stx1B targeted mice were viable until adulthood. We next performed Netrin-1 repulsion assays in EGL explants derived from P5 mice. EGL explants from control Stx1B (+/+) littermates displayed strong axonal chemorepulsion when confronted with Netrin-1 expressing cells. In contrast, chemorepulsion to Netrin-1 was significantly reduced in explants from homozygous Stx1B-deficient mice (**Figure 3C, D**), but was not completely abolished. To ascertain whether a functional redundancy with Stx1A may compensate for the lack of Stx1B, we generated double Stx1B/Stx1A-deficient mice. However, as double Stx1B/Stx1A knock-out animals die at birth (Ros *et al*., 2018), we used an alternative approach to target both Stx1 paralogs. We created a Stx1-shRNA sequence complementary to a conserved region in both Stx1A and 1B, enabling us to knock-down both Stx1 paralogs. The efficiency of Stx1 downregulation was confirmed in PC12 cells, in which expression of the shRNA Stx1A-1B construct decreased Stx1 protein levels detected by western blot and made Stx1 undetectable by immunocytochemistry (**Figure EV3C, D**). After electroporating EGL explants with the shRNA Stx1A-1B, the downregulation of both Stx1 paralogs resulted in a dramatic decrease in chemorepulsion, when compared to control cultures (**Figure 3E, F**). Taken together with the above results, these observations suggest that both Stx1 paralogs participate in Netrin-1-mediated chemorepulsion of EGL axons.

**Figure 3.**
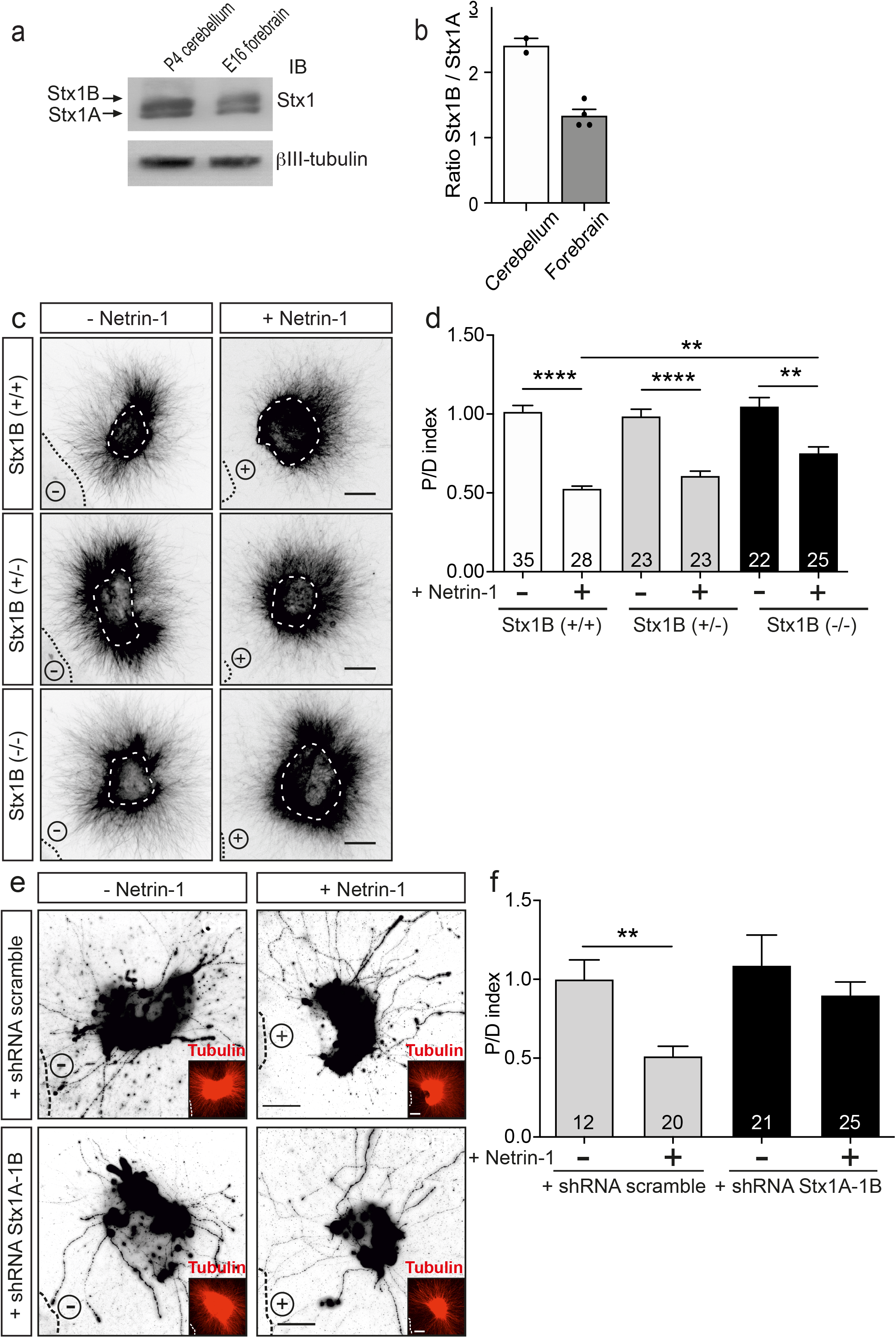
Downregulation of Stx1 blocks Netrin-1-induced repulsion in EGL explants. **(A)** Postnatal cerebellum and embryonic forebrain homogenates were subjected to urea/SDS-PAGE, resolving two Stx1 bands corresponding to Stx1B (upper band) and Stx1A (lower band). (**B)** Quantification of the ratio of intensities of Stx1B and Stx1A. (**C)** Representative images of EGL explants from P4 mice of different genetic backgrounds for Stx1B: wild-type Stx1B (+/+), heterozygous Stx1B (+/-), knock-out Stx1B (-/-). Explants were immunodetected with anti-βIII-tubulin. Explants are outlined with a white dashed line. (**D)** Graph showing the calculated P/D ratio. (**E)** Representative images from EGL explants from P4 mice. Explants were immunodetected with anti-GFP and anti-βIII-tubulin (inset images). Explants were electroporated with a shRNA scrambled control plasmid, or with a shRNA plasmid against both Stx1A and Stx1B (shRNA Stx1A-1B). (**F)** Graph showing the calculated P/D ratio. Explants in (**C)** and (**E)** were confronted with control HEK-293T cell aggregates (-Netrin-1) or Netrin-1-secreting aggregates (+ Netrin-1). Cell aggregates are outlined with a black dashed line in (**C)** and (**E)**. Scale bars in (**C)** and (**E)** represent 100 µm. Data in the plots in (**D)** and (**F)** represent mean ± SEM. One-way ANOVA followed by Dunn’s multiple comparison post hoc test of selected pairs was used. **p ≤ 0.01, ****p ≤ 0.0001. The number of explants analysed for each condition is indicated within each bar. Results were generated from at least three independent experiments.

### Netrin-1 induces growth cone collapse associated with membrane internalization

To better to understand the mechanisms involved in Netrin-1 chemorepulsion, we performed collapse assays in postnatal cerebellar EGL neurons. Primary neuronal cultures were exposed to Netrin-1 for 15-45 minutes and stained with phalloidin to label F-actin and analyse growth cone morphologies. Control cultures exhibited typical growth cones with triangular shapes and numerous lamellipodia and filopodia (**Figure 4A**). Incubation with Netrin-1 resulted in the collapse of up to 75% of growth cones (**Figure 4B**), which exhibited the typical shrivelled, round-type, pencil-like shape, devoid of filopodia or lamellipodia (**Figure 4A**).

**Figure 4.**
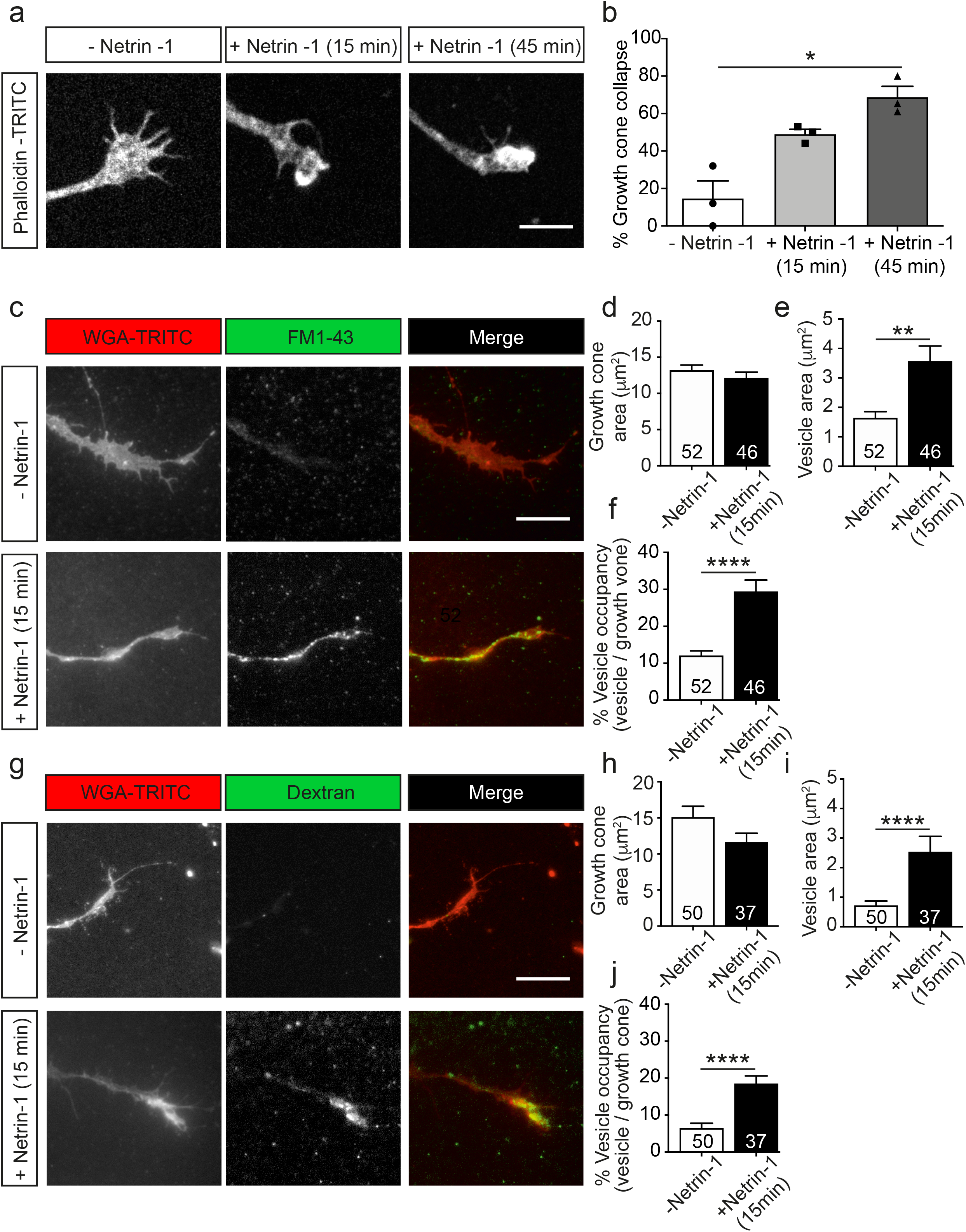
Netrin-1-induced growth cone collapse is associated with membrane internalization in EGL neurons. **(A)** Representative confocal images of growth cones from EGL neurons treated with either control medium or Netrin-1-supplemented medium (300 ng / mL, 15 min or 45 min). Neurons were then immunostained with phalloidin-TRITC to detect actin cytoskeleton. Scale bar represents 10 µm. (**B)** The percentage of collapsed growth cones was calculated and plotted for each time point. Data represent mean ± SEM. One-way ANOVA followed by Dunn’s multiple comparison post hoc test of selected pairs was used. *p ≤ 0.05. Each data point plotted represent one independent experiment. 51 to 55 growth cones were analysed per experiment. (**C)** Representative confocal images of growth cones from EGL neurons treated with either control medium or Netrin-1-supplemented medium (300 ng / mL), together with FM1-43, for 15 min. Neurons were then labelled with the plasma membrane marker WGA-TRITC. Scale bar represents 10 µm. (**D)** Growth cone area was measured and plotted for each condition. (**E)** The total area occupied by vesicles within the growth cones was measured and plotted for each condition. (**F)** The percentage of the growth cone occupied by endocytic vesicles was calculated and plotted for each condition. (**G)** Representative confocal images of growth cones from EGL neurons treated with either control medium or Netrin-1-supplemented medium (300 ng / mL), together with Alexa Fluor 488-LMW-dextran, for 15 min. Neurons were then labelled with the plasma membrane marker WGA-TRITC. Scale bar represents 10 µm. (**H)** Growth cone area was measured and plotted in a graph for each condition. (**I)** The total area occupied by vesicles within the growth cones was measured and plotted for each condition. (**J)** The percentage of the growth cone occupied by endocytic vesicles was calculated and plotted for each condition. Data in (**D**-**F)** and (**H**-**J)** represent mean ± SEM. Unpaired two-tailed Student’s *t* tests were used. **p ≤ 0.01, ****p ≤ 0.0001. The number of growth cones analysed for each condition is indicated within each bar. Obtained results were generated from at least three independent experiments.

Previous studies have shown that Semaphorin 3A (Fournier *et al*, 2000; Tojima *et al*., 2010) and Slit2 (Piper *et al*., 2006) induce endocytic events in growth cones. We therefore investigated whether Netrin-1 leads to endocytosis in EGL growth cones, using two typical markers of endocytosis, the styryl dye FM1-43 (Gaffield & Betz, 2007) and a fluorescently labelled low-molecular-weight (LMW, < 10 kDa) dextran (Alexa Fluor 488-LMW-dextran) (Fournier *et al*., 2000). After incubation with Netrin-1 and the above dyes, EGL cultures were incubated with wheat germ agglutinin conjugated with TRITC (WGA-TRITC) to label the cell surface of growth cones (**Figure 4C, G**). After 15 minutes of Netrin-1 treatment, the growth cone areas were slightly reduced, ranging from 8% to 23% reduction (**Figure 4D, H**). The total area of internalized vesicles within growth cones was also used to quantify the resulting endocytosis during growth cone collapse. Both markers used to monitor endocytosis showed that Netrin-1 triggers a marked increase in the area of endocytosed vesicles per growth cone (**Figure 4E, I**), as well as an increase in the percentage of the growth cone area occupied by endocytic vesicles (**Figure 4F, J**), suggesting that the growth cone starts collapsing through the internalization of its cell surface components.

### Macropinocytosis, but not clathrin-dependent endocytosis, mediates Netrin-1-induced growth cone collapse

To test whether Netrin-1-induced growth cone collapse requires clathrin-dependent endocytosis we used a combination of pharmacological and dominant negative strategies. Monodansylcadaverine (MDC) is a potent inhibitor of clathrin-dependent endocytosis (Schutze *et al*, 1999), and Tyrphostin A23 (AG18) is an inhibitor of the binding of internalized cargo to AP-2 (Banbury *et al*, 2003). We found that neither AG18 nor MDC was able to prevent the collapse of growth cones (**Figure 5A, B**), or the formation of Netrin-1-induced internalization of vesicles in growth cones (**Figure 5C**). Eps15-Δ95/295 is a dominant-negative form of Eps15 (EGFR pathway substrate clone 15) lacking the Eps15 homology domains. This truncated molecule inhibits the plasma membrane targeting of AP-2 and clathrin, as well as the formation of coated pits and clathrin-dependent endocytosis (Benmerah *et al*, 1999; Poupon *et al*, 2002). Mutant Eps15-Δ95/295 overexpression in EGL neurons did not prevent growth cone collapse in neurons treated with Netrin-1 (**Figure 5D, E**). These results suggested that Netrin-1-induced large vesicle endocytosis and growth cone collapse are independent of clathrin-dependent internalization mechanisms.

**Figure 5.**
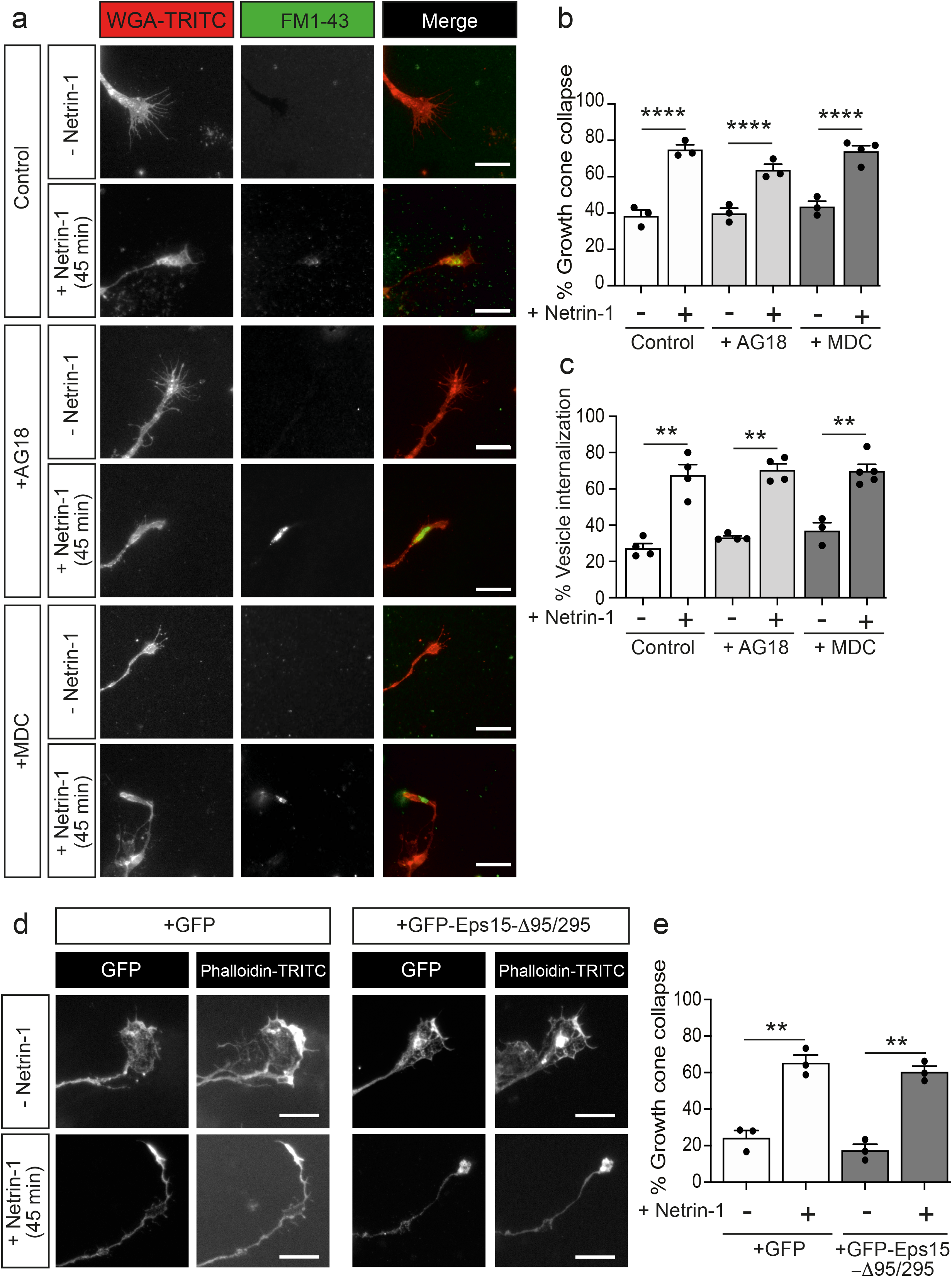
Netrin-1-induced growth cone collapse is independent of clathrin-dependent endocytosis in EGL neurons. (**A)** Representative confocal images of growth cones from EGL neurons treated with either control medium or Netrin-1-supplemented medium (300 ng / mL), together with FM1-43, for 45 min. 30 min before the addition of control medium or Netrin-1, neurons were supplemented with control (DMSO) or with the clathrin-dependent internalization inhibitors MDC (1 μM), or AG18 (100 μM). These reagents were maintained during the subsequent 45 min incubation with Netrin-1. Neurons were then labelled with the plasma membrane marker WGA-TRITC. Scale bar represents 10 µm. (**B)** The percentage of collapsed growth cones was calculated and plotted for each condition. (**C)** The percentage of growth cones with internalized vesicles was plotted for each treatment. (**D)** Representative confocal images of growth cones from EGL neurons transfected with GFP or with the dominant negative GFP-Eps15-Δ95/295, and treated with either control medium or Netrin-1-supplemented medium (300 ng / mL, 15 min or 45 min). Neurons were then immunostained with phalloidin-TRITC to detect actin cytoskeleton. Scale bar represents 10 µm. (**E)** The percentage of collapsed growth cones was plotted for each condition. Data of the plots in (**B)**, (**C)** and (**E)** represent mean ± SEM. One-way ANOVA followed by Dunnett’s multiple comparison post hoc test of selected pairs was used. **p ≤ 0.01, ****p ≤ 0.0001. Each data point represents one independent experiment. From 27 to 129 growth cones were analysed per experiment.

Macropinocytosis is a clathrin-independent form of endocytosis that leads to the formation of large endocytic vacuoles (Ferreira & Boucrot, 2018). Incorporation of high-molecular-weight (HMW, > 10 kDa) dextran has been associated with membrane retrieval by macroendocytic vesicles (Kabayama *et al*., 2009; Kolpak *et al*, 2009). We investigated whether Netrin-1 induced membrane retrieval through macropynocytosis by analysing the internalization of Fluorescein-HMW-dextran in the absence or presence of 5-(N-ethyl-N-isopropyl) amiloride (EIPA), a potent analogue of amiloride channels that has been used as a macropinocytosis-specific inhibitor (Meier *et al*, 2002). Our results revealed that incubation with Netrin-1 induced the internalization of HMW-dextran, but this internalization was completely blocked by pre-treatment with EIPA (**Figure 6A, B**). We next asked whether the blockade of macropynocytosis also influenced Netrin-1-induced growth cone collapse. Pre-incubation with EIPA was able to dramatically abolish Netrin-1-induced growth cone collapse in EGL growth cones (**Figure 6C, D**). Taken together, our data indicate that macropinocytosis, but not clathrin-dependent endocytosis, mediates Netrin-1-induced growth cone collapse in EGL axons.

**Figure 6.**
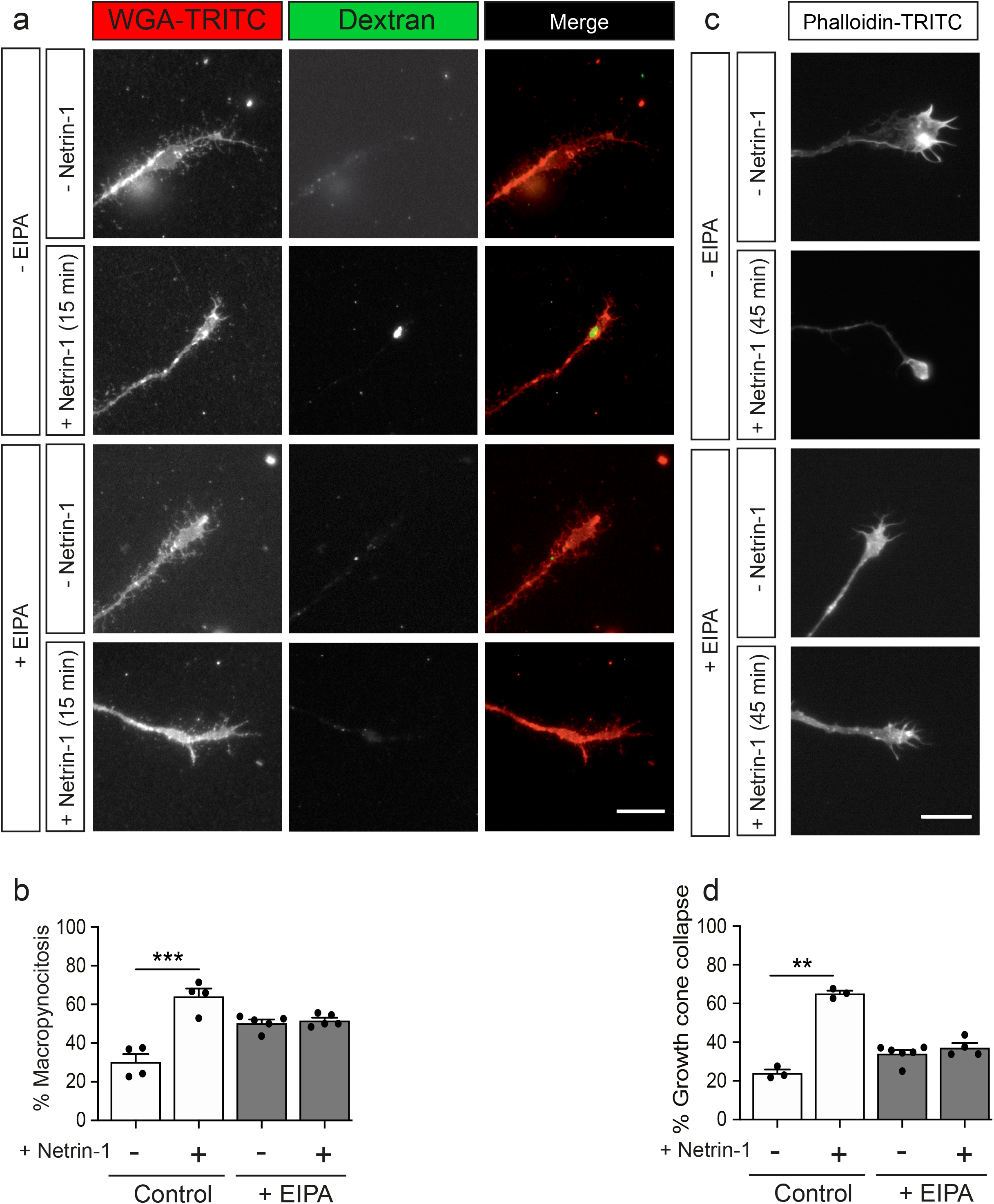
Netrin-1-induced growth cone collapse is dependent of macropinocytosis in EGL neurons. (**A)** Representative confocal images of growth cones from EGL neurons treated with either control medium or Netrin-1-supplemented medium (300 ng / mL), together with Fluorescein-HMW-dextran, for 15 min. 10 min before the addition of control medium or Netrin-1, neurons were supplemented with control (DMSO) or with the macropinocytosis-specific inhibitor EIPA (10 μM). They were maintained on these supplements for the subsequent 15 min incubation with Netrin-1. Neurons were then labelled with the plasma membrane marker WGA-TRITC. Scale bar represents 10 µm. (**B)** The percentage of growth cones with macropinocytic vesicles plotted for each treatment. (**C)** Representative confocal images of growth cones from EGL neurons treated with either control medium or Netrin-1-supplemented medium (300 ng / mL) for 45 min. 10 min before the addition of control medium or Netrin-1, neurons were supplemented with control (DMSO) or with the macropinocytosis-specific inhibitor EIPA (10 μM), which was maintained during the subsequent 45 min incubation with Netrin-1. Neurons were then immunostained with phalloidin-TRITC to detect actin cytoskeleton. (**D)** The percentage of collapsed growth cones was plotted for each treatment. Data in the plots in (**B)** and (**D)** represent mean ± SEM. One-way ANOVA followed by Dunn’s multiple comparison post hoc test of selected pairs was used. **p ≤ 0.01, ***p ≤ 0.001. Each data point represents one independent experiment. 55 to 185 growth cones were analysed per experiment.

### Stx1 blockade inhibits Netrin-1-induced growth cone collapse and macropinocytosis

It has recently been reported that Stx1 is involved in rapid and slow endocytosis at synapses, which can be either clathrin-independent or -dependent (Xu *et al*., 2013). As Stx1 interacts with UNC5 receptors, which mediate chemorepulsion and growth cone collapse, we next determined whether Netrin-1-induced growth cone macropynocitosis and collapse was Stx1-dependent using BoNTs. BoNT/A or BoNT/C1 was added to the medium 10 minutes before incubation with Netrin-1. The percentage of macropinocytic vesicle-containing growth cones increased about two fold when Netrin-1 was added alone or in the presence of BoNT/A (**Figure 7A, C)**. In contrast, Netrin-1-induced formation of macropinocytic vesicles was specifically reduced by cleaving Stx1 using BoNT/C1 (**Figure 7A, C**). We then tested whether BoNT/A or BoNT/C1 altered growth cone collapse. Neurons treated with control medium or with control medium supplemented with BoNT/A or BoNT/C1 exhibited normal growth cone morphologies with lamellipodia and filopodia (**Figure 7B**). After incubation with Netrin-1, about 70% of EGL growth cones exhibited collapsed and round-tipped shapes (**Figure 7B, D**). This effect was maintained when EGL neurons were co-incubated with BoNT/A, but was markedly reduced when the cultures were co-incubated with BoNT/C1 (**Figure 7D**). These results suggested that Stx1 is necessary for both Netrin-1-induced growth cone collapse and macropinocytosis.

**Figure 7.**
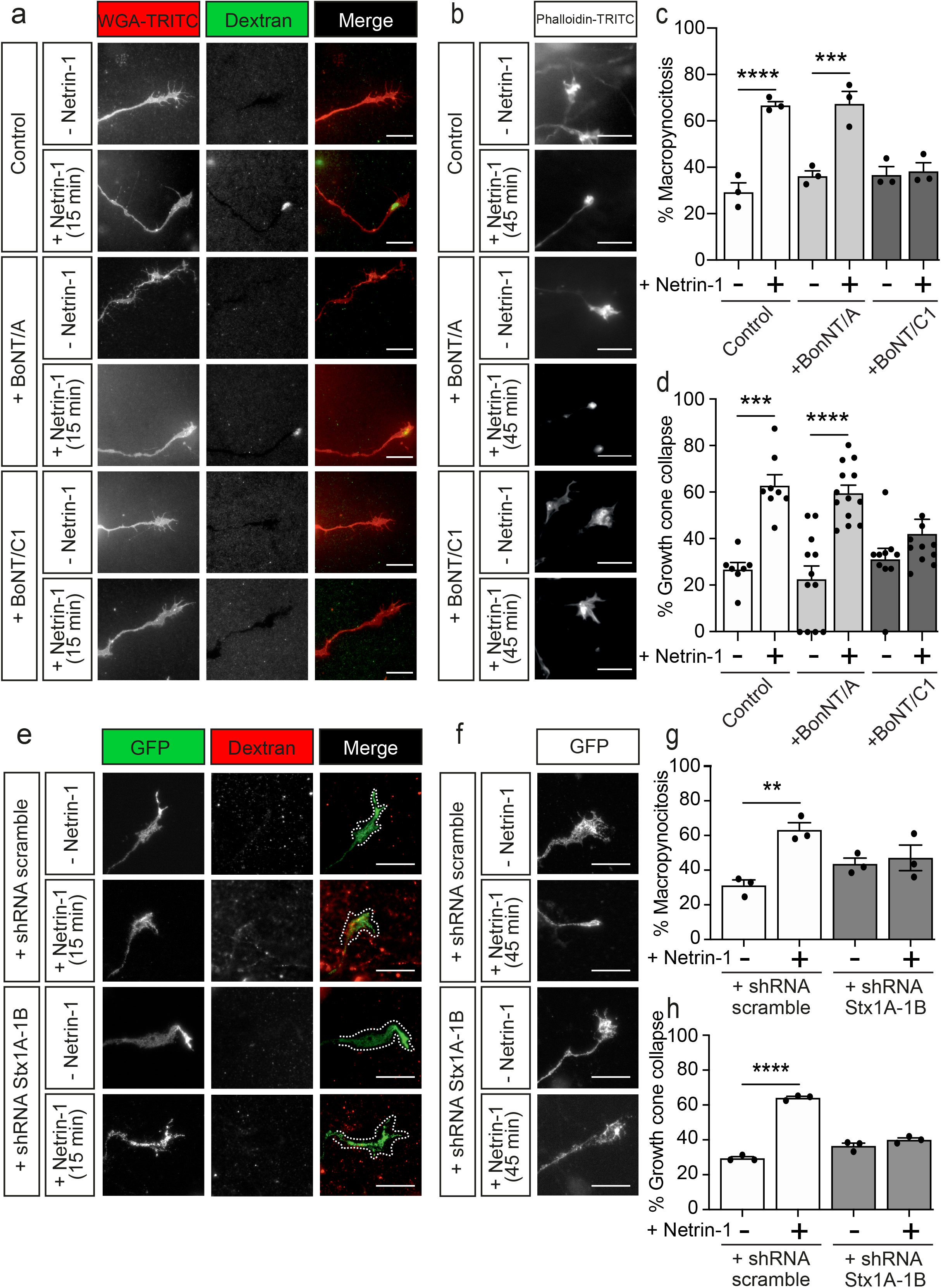
Netrin-1-induced growth cone collapse requires Stx1 in EGL neurons. (**A)** Representative confocal images of growth cones from EGL neurons treated with either control medium or Netrin-1-supplemented medium (300 ng / mL), together with Fluorescein-HMW-dextran, for 15 min. 15 min before the addition of control medium or Netrin-1, neurons were supplemented with control (PBS) or 25 nM BoNT/A (+ BoNT/A) or 15 nM BoNT/C1 (BoNT/C1). The toxins were maintained during the subsequent 15 min incubation with Netrin-1. Neurons were then labelled with the plasma membrane marker WGA-TRITC. Scale bar represents 10 µm. (**B)** Representative confocal images of growth cones from EGL neurons treated with either control medium or Netrin-1-supplemented medium (300 ng / mL). 15 min before the addition of control medium or Netrin-1, neurons were supplemented with control (PBS) or 25 nM BoNT/A (+ BoNT/A) or 15 nM BoNT/C1 (BoNT/C1). The toxins were maintained during the subsequent 45 min incubation with Netrin-1. Neurons were then immunostained with phalloidin-TRITC to detect actin cytoskeleton. Scale bar represents 10 µm. (**C)** The percentage of growth cones with macropinocytic vesicles was calculated and plotted in a graph for each treatment. (**D)** The percentage of collapsed growth cones was plotted for each treatment. (**E)** Representative confocal images of growth cones from EGL neurons transfected with a shRNA scrambled control plasmid, or with a shRNA plasmid against both Stx1A and Stx1B (shRNA Stx1A-1B), and treated with either control medium or Netrin-1-supplemented medium (300 ng / mL), together with tetramethylrhodamine-HMW-dextran, for 15 min. Neurons were then immunostained anti-GFP to detect transfected growth cones. Scale bar represents 10 µm. (**F)** Representative confocal images of growth cones from EGL neurons transfected with a shRNA scrambled control plasmid, or with a shRNA plasmid against both Stx1A and Stx1B (shRNA Stx1A-1B), and treated with either control medium or Netrin-1-supplemented medium (300 ng / mL), for 45 min. Neurons were then immunostained with anti-GFP to detect transfected growth cones. Scale bar represents 10 µm. (**G)** The percentage of growth cones with macropinocytic vesicles was plotted for each treatment. (**H)** The percentage of collapsed growth cones was plotted for each treatment. Data of the plots in (**C)**, (**D)**, (**G)** and (**H)** represent mean ± SEM. One-way ANOVA followed by Sidak’s multiple comparison post hoc test of selected pairs was used. **p ≤ 0.01, ***p ≤ 0.001, ****p ≤ 0.0001. Each data point represents one independent experiment. 7 to 68 growth cones were analysed per experiment.

To confirm these findings, we downregulated both Stx1 paralogs using the Stx1A-1B shRNA construct. Electroporation of EGL cells with Stx1A-1B shRNA, but not with a control scrambled shRNA sequence, blocked the formation of macropinocytic vesicles in EGL growth cones incubated with Netrin-1 (**Figure 7E, G**). Similarly, the knockdown of Stx1 blocked Netrin-1-dependent collapse of growth cones (**Figure 7F, H**). Taken together, our results therefore demonstrate that Stx1 is required for both Netrin-1-dependent growth cone collapse and micropinocytosis (**Figure 8**).

**Figure 8.**
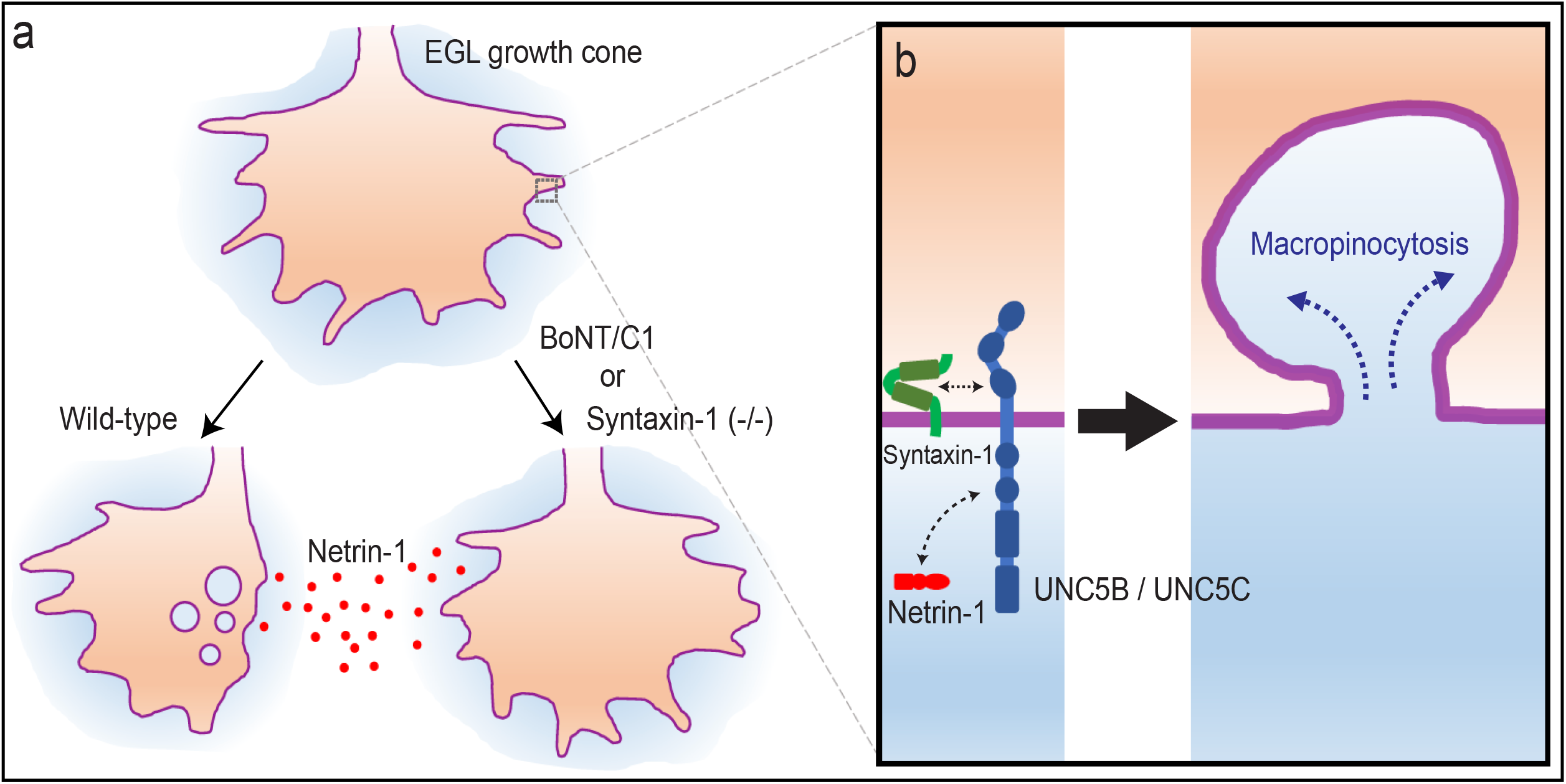
Model of Netrin-1-dependent chemorepulsion in EGL growth cones. **(A)** In wild-type conditions, Netrin-1 induces chemorepulsion of postnatal EGL axons through a mechanism that initiates the formation of clathrin-independent large micropinocytic vesicles in the growth cone. This retrieval of plasma membrane is the mechanism responsible for the collapse of EGL growth cones. However, either the absence of syntaxin-1 or its inactivation using specific toxins (BoNT/C1) prevents both the micropinocytosis and the subsequent collapse of EGL growth cones. **(B)** Detail of the molecular events during Netrin-1-mediated growth cone collapse. Syntaxi-1 interacts with UNC5 receptors in EGL neurons (UNC5B and UNC5C). The binding of Netrin-1 initiates the formation of large macropinocytic vesicles.

## Discussion

The dynamics of growth cones is essential for axonal guidance and the establishment of neuronal connectivity. How the growth cone transduces guidance cue signalling into cellular responses has received notable attention. The motile nature of the growth cone requires fine control of cytoskeletal and membrane dynamics (Blanquie & Bradke, 2018; McCormick & Gupton, 2020). The growth cone is known to be a major site of endo- and exocytosis, but the mechanisms that regulate these processes in response to chemoattractive or chemorepulsive cues are still being unveiled. Based on the results of the current study, we propose a new mechanism involving the close interplay between signalling receptors and membrane homeostasis by means of the SNARE protein Stx1. We show that a biochemical interaction occurs between the repulsive Netrin-1 receptor UNC5 and the SNARE protein Stx1. We also demonstrate that macropinocytosis, and not clathrin-dependent endocytosis, is the membrane retrieval process mediating repulsive responses to Netrin-1, and that the absence of Stx1 or interference with its function blocks repellent Netrin-1 responses in EGL cells, from membrane endocytosis to chemorepulsion.

Two major mechanisms of membrane retrieval have been associated with axon guidance: clathrin-dependent endocytosis and macropincytosis (Kabayama *et al*., 2009; Kolpak *et al*., 2009; Tojima *et al*., 2010). We investigated growth cone endocytosis through two complementary approaches, using staining of newly formed vesicles with FM dyes or with fluorescent dextran. FM dyes have previously been used to study growth cone endocytosis, revealing a wide plethora of vesicular structures with different morphologies and sizes (Hines *et al*., 2010), and the incorporation of HMW dextrans has been associated with macroendocytic events (Kabayama *et al*., 2009; Kolpak *et al*., 2009; Swanson & Watts, 1995). Using both markers, FM dyes stained virtually all dextran-positive vesicles whereas the dextran-positive vesicles account for less than half of those stained with FM dyes (Clayton & Cousin, 2009). Our data reproduce this phenomenon and are consistent with the existence of at least two vesicle populations that are regulated by distinct signalling cascades, with a pool of large vesicles generated by macropinocytosis through the repulsion by Netrin-1. Recent studies have described the mechanisms underlying endocytic processes in response to class III semaphorins, but the characterization of membrane dynamics during repulsive responses to Netrin-1 has yet to be addressed (Boyer & Gupton, 2018; Fournier *et al*., 2000; Itofusa *et al*, 2017; Jurney *et al*., 2002). In cortical growth cones, the repulsive actions of Semaphorin-3A have been hypothesized to require a VAMP-2-dependent mechanism of receptor sorting (Zylbersztejn *et al*., 2012) and clathrin-dependent endocytosis of local membrane. Our results suggest that the growth cone collapse and retraction associated with Netrin-1 are not affected by inhibition of clathrin-dependent coated pit formation of endocytic vesicles, and are therefore not involved in Netrin-1-induced repulsion of EGL axons. On the other hand, macropinocytosis is considered the main cellular mechanism to uptake large volumes of liquid and membrane. Macropinosomes can be generated by actin-dependent membrane ruffling or by the fusion of multiple membrane ruffles with the plasma membrane, allowing the incorporation of HMW molecules such as dextrans (Swanson & Watts, 1995). Our data show an association of growth cone collapse with an increase on large, dextran-positive vesicle incorporation upon Netrin-1 treatment. This increase was abolished in the presence of the macropinocytosis-specific inhibitor EIPA (Kabayama *et al*., 2009; Meier *et al*., 2002), together with the suppression of Netrin-1-induced growth cone collapse. These findings demonstrate for the first time that Netrin-1-induced repulsion relies on macropinocytosis as the mechanism for growth cone membrane retrieval, in contrast to repulsion to class-III semaphorins, and analogous to Shh repulsion (Kolpak *et al*., 2009). Our study therefore confirms that repulsive guidance relies on multiple membrane retrieval mechanisms whereby a given set of growth-inhibiting molecules associate with different signalling pathways to drive endocytosis.

Netrin-1 was the first guidance cue to be described but the intricacies of its functions are still being revealed (Boyer & Gupton, 2018). Netrin-1 exerts a strong role during cerebellar development, controlling granule cell migration, and regulating the formation of contralateral projections towards the midline, the formation of ipsilateral projections during the extension of parallel fibres, and the development of olivo-cerebellar projections (Bloch-Gallego *et al*, 1999). Proteins involved in the downstream signalling cascades associated with growth cone repulsion also produce alterations in granule cell migration from the EGL and unresponsiveness to Netrin-1 (Peng *et al*, 2010). These results suggest a predominant role of Netrin-1 in correct cerebellar formation. Our results support these studies and offer a new view of this mechanism, whereby the induction of macropinocytosis underlines the repulsive effects of Netrin-1.

SNARE proteins are expressed in the developing brain and axons, and their role beyond synaptic transmission during axon navigation is just being unveiled. VAMP-2 function is required for repulsion to class III semaphorins (Zylbersztejn *et al*., 2012). We have previously described how Stx1 mediates ligand-dependent exocytosis and attraction to neurotrophic factors (Fuschini *et al*., 2018) and Netrin-1 (Cotrufo *et al*., 2012; Cotrufo *et al*., 2011) through the interaction with Trk receptors and DCC, respectively. Here we describe how Stx1 interacts directly with the repulsive Netrin-1 receptor UNC5. The repulsive guidance of EGL axons was severely impaired when interfering with Stx1 function through i) its cleaving with BoNT/C1, ii) its suppression in knock-out animals or iii) its downregulation using specific shRNAs. Both Stx1A and 1B are abundantly expressed in the brain (Aguado *et al*, 1999) and share basic functions as neuronal t-SNAREs. However, only Stx1B is necessary for the regulation of spontaneous and evoked synaptic vesicle exocytosis in fast transmission (Mishima *et al*, 2014), which suggests that compensatory mechanisms underlie the mild impairment in basal synaptic transmission reported in Stx1A knock-out animals (Fujiwara *et al*., 2006). In addition, Stx1B is essential for spontaneous GABAergic transmission frequency in the cerebellum, most likely attributed to a lower number of neurons, suggesting a differential importance of Stx1B for neuronal survival (Park *et al*, 2014; Wu *et al*, 2015). Our approach using Stx1B knock-out mice only partially reduced the repulsive effects of Netrin-1 on EGL neurons. Experiments with Stx1B (+/+), (+/-), and (-/-) animals showed an incremental dose-dependence in the severity of their phenotypes, appointing to a partial functional compensation. Additional experiments in double knock-out animals may have confirmed this compensatory effect, but the embryonic lethality of these mice precluded such studies.

Our work demonstrates for the first time that Stx1 is required for Netrin-1-induced macropinocytosis and growth cone collapse. It has previously been shown that Stx1 and other SNARE proteins also play a role in endocytosis at synapses (Xu *et al*., 2013; Zhang *et al*., 2013), but no mechanistic explanation has been proposed. Our experiments using toxins further indicate that Stx1 function, and not synaptic transmission mediated by the SNARE complex, is required to transduce Netrin-1 chemorepulsion, given that BoNT/A treatments, which specifically disrupt SNAP-25 but do not affect Stx1, do not block repulsion to Netrin-1. In addition, they provide mechanistic insight, as interfering with Stx1 function using BoNT/C1 or shRNA also blocks the Netrin-1-induced increase in macropinocytosis that precedes growth cone collapse. These results cannot be explained by previous reports that Stx1 downregulation produces an enhancement of basal growth cone collapse and macropinocytosis (Igarashi *et al*, 1996; Kabayama *et al*., 2009). The authors’ hypothesis of an imbalance in newly exocytosed membrane retrieval due to the absence in Stx1 is supported by our findings, but fails to completely explain the blockade of macropinocytosis and collapse upon Netrin-1 treatments that we observed. In previous reports we demonstrate that Stx1 forms a protein complex with the Netrin-1 receptor DCC, coupling the chemotropic Netrin-1/DCC axonal attraction and growth cone SNARE-dependent exocytic fusion of vesicles with extracellular membranes (Cotrufo *et al*., 2011). In agreement with these findings, here we demonstrated that the t-SNARE Stx1 is also able to interact physically with the Netrin-1 receptor UNC5, thereby mediating cerebellar repulsion.

Binding studies have found that Stx1 interacts with Dynamin-2 during secretion in adrenal chromaffin cells (Galas *et al*., 2000). It has been reported that Dynamin-2 is involved in clathrin-dependent endocytosis (Bertot *et al*, 2018) and in clathrin-independent macropinocytosis induced by Shh (Kolpak *et al*., 2009). Thus, Dynamin-2 could be mediating the macropinocytic response to Netrin-1 in co-operation with Stx1 and UNC5 receptors. A parallel role for Stx1 could reside in the formation of the pseudopodial extension that precedes phagosome formation, as it requires focal exocytosis and reorganization of the actin cytoskeleton (Lim & Gleeson, 2011). The mechanisms required for focal exocytosis in macrophage macropinocytosis resemble those involved in neuronal growth cone extension, as both require SNARE proteins such as Ti-VAMP (Braun *et al*, 2004; Cotrufo *et al*., 2011) to provide the new membrane that is necessary for extension. Additional experimentation to investigate the mechanistic machinery involving Stx1 during Netrin-1/UNC5-induced growth cone chemorepulsion is warranted.

In conclusion, attractive and repulsive responses to Netrin-1 use a parallel approach to handle membrane incorporation and retrieval. Both DCC and UNC5 rely on Stx1 to initiate membrane exocytosis or endocytosis and ultimately attractive or repulsive growth cone responses. This opens a new scenario whereby key proteins share multiple apparently opposite roles depending on the context of the signalling molecules present.

## Matierials and Methods

### Plasmids

Full-length Stx1A and Stx1A-GFP were created as described previously (Cotrufo *et al*., 2011). UNC5A-myc, UNC5B-myc and UNC5C-myc were a gift from Prof. Lindsay Hinck (University of California, Santa Cruz), pEGFP-C1 (Clontech), pcDNA3.1 (Invitrogen, Thermo Fisher Scientific), GFP-Eps Δ95/295 vector was a gift from Dr. Francesc Tebar (University of Barcelona, Spain). For shRNA experiments, the pLVTHM plasmid was kindly provided by Prof. Didier Trono (École polytechnique fédérale de Lausanne, Lausanne), including specific oligonucleotides for Stx1A and Stx1B sequence: gatccccCCAGAGGCAGCTGGAGATCACttcaagagaGGTCTCCGTCGACCTCTAGT Gttttt (Forward), and agctaaaaaCCAGAGGCAGCTGGAGATCACtctcttgaaGGTCTCCGTCGACCTCTAGT Gggg (Reverse).

### Heterologous cell cultures

HEK-293T cells were maintained in DMEM (11995065; GIBCO-Thermo Fisher Scientific) medium supplemented with 10% fetal bovine serum (FBS; 26140079; GIBCO-Thermo Fisher Scientific), 1% GlutaMAX (35050061; GIBCO-Thermo Fisher Scientific) and 1% penicillin/streptomycin (15140122; GIBCO-Thermo Fisher Scientific). PC12 cells were maintained in DMEM containing 1% GlutaMAX, 5% FBS, 5% horse serum (HS; 26050-088; GIBCO-Thermo Fisher Scientific), and 1% penicillin/streptomycin.

### Primary neuronal cultures

Primary cultures and explants of cerebellar EGL neurons were prepared from P3-P5 postnatal CD1 strain mice (Charles River). Animals were sacrificed by decapitation in accordance with institutional and governmental ethical guidelines and regulations. All the experiments using animals were performed in accordance with the European Community Council directive and the National Institutes of Health guidelines for the care and use of laboratory animals. Experiments were also approved by the ethical committee from the Generalitat of Catalonia. Postnatal cerebellums were isolated, mechanically disaggregated and trypsinized as previously described (Hernaiz-Llorens *et al*., 2020; Rosello-Busquets *et al*, 2019). Briefly, after centrifugation, neurons were resuspended in 2 mL of DMEM medium and EGL neurons were isolated by centrifugation (3000 rpm, 10 min at 4° C) in a discontinuous percoll gradient (35% and 60% of percoll). After washing with PBS, EGL neurons were plated on poly-D-lysine pre-coated dishes in DMEM, 1% penicillin/streptomycin, 1% glutamine, 4.5% D-(+)-glucose (G-8769; Sigma), 5% HS and 10% FBS. This medium was maintained for 24 h and then changed to a medium where HS and FBS were replaced by 2% B27 (17504001; GIBCO-Thermo Fisher Scientific) and 1% N2 (11520536; GIBCO-Thermo Fisher Scientific). Neurons were cultured for 72 h (3DIV).

To isolate and culture EGL explants, cerebellums from P3-P5 mice were isolated and chopped in to 300 µm slices. Selected slices were further dissected using fine tungsten needles to extract small tissue pieces from the EGL. Explants were carefully placed inside a 3D collagen matrix on poly-D-lysine pre-coated dishes, prepared as previously described(Gil & Del Rio, 2019). Explants were cultured for 48 h (2DIV) in DMEM, 1% penicillin/streptomycin, 1% glutamine, 4.5% D-(+)-glucose, 2% B27.

### Transfection of heterologous cells and primary neurons, and electroporation of explants

One day prior to transfection, cells were counted and plated in 100 mm dishes, to have a 70% of confluence on the day of transfection. HEK-293T or PC12 cells were transfected with Lipofectamine 2000 (11668019; Thermo Fisher Scientific) following the manufacturer’s instructions.

After 2DIV, neuronal cultures were transfected using Lipofectamine 2000. Prior to transfection, 3/5 of medium was removed and kept aside to be returned later. For each 35 mm plate, 4 µg of DNA and 8 µL of Lipofectamine 2000 were used. The final mixture of DNA-Lipofectamine 2000 was carefully added to the cultures and further incubated at 37° C in 5% CO_2_ for 60 min. The transfection medium containing DNA and Lipofectamine 2000 was finally replaced by the medium removed at the beginning. An equal amount of freshly prepared medium was added. Cultures were incubated until the following day.

Explants were electroporated using the Invitrogen Neon system. Briefly, dissected explants were washed three times in PBS and resuspended in buffer R containing 2 to 4 µg of the DNA to electroporate. The conditions for the electroporation were voltage: 500 V; width: 50 ms; 5 pulses. Explants were immediately washed once in Neurobasal medium and then mounted in a 3D collagen matrix.

### Immunocytochemistry

PC12 cells and primary neurons were fixed with a solution of 4% paraformaldehyde (PFA) in PBS for 10 min at room temperature. Neuronal explants were fixed by incubation with the same solution for 30 min at room temperature. They were rinsed with PBS, and permeabilized with a solution of 0.1% Triton-X-100 in PBS for 10 min, or 30 min for the explants. Cells were then washed with PBS and incubated in blocking solution (10% HS in PBS) for 1 h at room temperature, or 3 h for the explants. After blocking, the cells were incubated with the respective primary antibodies diluted in blocking solution for 2 h at room temperature, or overnight at 4° C for the explants. Cells and explants were washed with PBS and incubated in the secondary antibody solution (PBS 1x with 10% HS) for 1 h at room temperature, or 3 h for the explants. Finally, cells and explants were washed and mounted in Mowiol (81381; Sigma-Aldrich) for imaging.

### Growth cone collapse and vesicle internalization experiments

EGL primary neurones were cultured for 3DIV and then treated with specific reagents or toxins, or the respective controls (DMSO or PBS), which were included 30 to 10 min before the addition of Netrin-1, and maintained during the whole incubation. In growth cone collapse experiments, neurons were then incubated with Netrin-1 (300 ng / mL) or control (BSA 0.1%) for 45 min at 37°C in DMEM containing 1% glutamine and 4.5% D-(+)-glucose. After this incubation, neurons were fixed with 4% PFA and permeabilized with PBS-Triton-X-100 (0.1%). Finally, actin filaments were stained by incubation with phalloidin-TRITC (1 µM) for 30 min. Cells were mounted on Mowiol and used for imaging. Actin staining was used to identify growth cones, which were outlined based on differential staining for actin in this compartment with respect to the adjacent axon. An intensity threshold mask was created using Fiji (Schneider *et al*, 2012) and the growth cone perimeter was selected using the wand tool. Collapsed growth cones were manually identified based on their morphology. In contrast to normal extended growth cones, collapsed growth cones lose their morphology, acquiring a shrivelled, round-tipped pencil-like shape devoid of lamellae or filopodia.

### Repulsion experiments

During the plating procedure, explants embedded into the 3D collagen matrix were confronted at a distance of 200 -600 µm with aggregates of HEK-293T cells stably expressing Netrin-1 (Cotrufo *et al*., 2011), or transiently transfected with semaphorin 3A or semaphorin 3F. Netrin-1 expression in stable cells was regularly checked by western blot (data not shown). If required, explants were treated with specific BoNTs (25 nM BoNT/A or 15 nM BoNT/C1) by adding them to the medium 3 to 4 h after being cultured. After 48 h (2DIV) of culture, explants were fixed, and immunocytochemistry against βIII-tubulin was performed. Repulsion was analysed by measuring the proximal/distal (P/D) ratio as previously described (Gil & Del Rio, 2019; Hernaiz-Llorens *et al*., 2020). Briefly, the P/D ratio is the fraction of axons growing in the proximal quadrant of the explant (closer to the aggregate of HEK-293T cells) versus the axons growing in the distal quadrant. A ratio close to 1 indicates a radial pattern of growth, below 1 indicates chemorepulsion and above 1 indicates chemoattraction.

### Statistics

Results were analysed statistically using GraphPad Prism software (GraphPad Software, Inc). The D’Agostino and Pearson test was used to test for normality. The unpaired two-tailed Student’s t test was used for comparison of two groups. For datasets comparing more than two groups, ANOVA followed by the Dunn test or the Dunnet test corrections for multiple comparison was used. Statistical comparisons were performed on a per-cell, per-explant or per-experiment basis. The neurons were randomly selected within the dishes and blindly analysed, and were collected from at least three independent experiments. Values are represented as the mean ± SEM. The tests used are indicated in the respective figure legends. If the number of elements analysed for each condition was above 15, it is indicated within each bar. If the number was below 15, each individual value is plotted. A p-value below 0.05 was accepted as significant.

### Ethical standards

All the experiments using animals were performed in accordance with the European Community Council directive and the National Institutes of Health guidelines for the care and use of laboratory animals. Experiments were also approved by the ethical committee from the Generalitat of Catalonia.

## Acknowledgements

We thank Julien Collombelli and the scientific staff of the Institute for Research in Biomedicine Microscopy Facility, for expert help with microscopy and data analysis, Alba del Valle for technical support and Rowan Tweedale and Alessandra Donato from QBI for critical reading and editing of the manuscript. This work was supported by grants from the Spanish MINECO (SAF2016-76340R and PID2019-106764RB-C21), CIBERNED (ISCIII), Spanish MECD (FPU14/02156, Excellence Unit María de Maeztu/Institute of Neurosciences, and BES-2014-067857), TERCEL (RD12/0019/0011), ERDF funds and by the “ Generalitat de Catalunya” (2014SGR1609). R.M-M was supported by the Juan de la Cierva postdoctoral fellowship. M.H-Ll was supported by MECD (BES-2014-067857). C.R-B was supported by MECD (FPU14/02156). T.C is supported through a Serra Hunter Fellow.

## Author contributions

R.M-M., A.M. and E.S. designed the research. R.M-M., A.M., C.R-B., T.C., M.H-Ll., R.M.A., F.P.B. and M.P. performed the experiments and analysed the data. R.M-M., O.R. and E.S. wrote the paper. R.M-M. and E.S. supervised the study.

## Expanded View Figure Legends

**Figure EV1. Stx1 does not interact with Semaphorin receptors. (A)** HEK-293T cells were transfected with the indicated combination of plasmids (empty vector pcDNA3, NP1-HA, NP1-HA and Stx1A, NP1-HA and Stx1A-GFP, PlexinA1-VSV, PlexinA1-VSV and Stx1A, or PlexinA1-VSV and Stx1A-GFP). Protein lysates were immunoprecipitated (IP) with anti-GFP, anti-HA or anti-VSV antibodies. Co-immunoprecipitation was detected by immunoblotting (IB) using anti-GFP (to detect Stx1A), anti-HA (to detect NP1) or anti-VSV (to detect PlexinA1) antibodies. **(B)** HEK-293T cells were transfected with the indicated combination of plasmids (empty vector pcDNA3, NP1-HA, NP1-HA and Stx1A, NP1-HA and Stx1A-GFP, PlexinA1-VSV, PlexinA1-VSV and Stx1A, or PlexinA1-VSV and Stx1A-GFP). Protein lysates were immunoprecipitated with anti-Stx1, anti-HA or anti-VSV antibodies. Co-immunoprecipitation was detected by immunoblotting using anti-Stx1 (to detect Stx1A), anti-HA (to detect NP1) or anti-VSV (to detect PlexinA1) antibodies.

**Figure EV2. (A)** Lysates from untreated EGL neurons or those treated with BoNT/C1 (15 nM) or BoNT/A (25 nM) for 45 min were immunoblotted against anti-Stx1, anti-SNAP-25 or anti-βIII-tubulin. **(B)** Representative images of hippocampal explants from E16 mice, immunodetected with anti-βIII-tubulin. Explants were confronted with Semaphorin 3A-or Semaphorin 3F-secreting aggregates. Explants were cultured in the absence of BoNTs (control) or in medium supplemented with 25 nM BoNT/A (+ BoNT/A) or 15 nM BoNT/C1 (BoNT/C1). HEK-293T aggregates are outlined with a dashed line. Scale bars represent 100 µm.

**Figure EV3. (A)** Western blot of embryonic forebrain homogenates from wild-type Stx1B (+/+), heterozygous Stx1B (+/-) and knock-out Stx1B (-/-) mice were subjected to urea/SDS-PAGE, resolving two Stx1 bands corresponding to Stx1B (upper band) and Stx1A (lower band). **(B)** Representative images of brains from embryonic E17 mice with different genetic backgrounds (Stx1B (+/+) -cyan-, Stx1B (+/-) -magenta-and Stx1B (-/-) -green-), together with their overlapping silhouettes to compare their size. **(C)** Western blot of lysates from PC12 cells transfected with a shRNA scrambled control plasmid, or with a shRNA plasmid against both Stx1A and Stx1B (shRNA Stx1A-1B). Samples were subjected to urea/SDS-PAGE to identify two Stx1 bands corresponding to Stx1B (upper band) and Stx1A (lower band). **(D)** Representative images of PC12 cells transfected with a shRNA scrambled control plasmid, or with a shRNA plasmid against both Stx1A and Stx1B (shRNA Stx1A-1B), and immunostained with anti-GFP to detect transfected cells (shRNA), with anti-Stx1 and with DAPI. Note that Stx1A-1B shRNA-transfected cells (arrowheads) display no Stx1 immunolabelling. Scale bar represents 10 µm.

